# The ultrafine-bridge-associated endonuclease ANKLE1 is stimulated by tension in DNA to process branch-points

**DOI:** 10.1101/2025.07.17.664873

**Authors:** Korak Kumar Ray, Artur P. Kaczmarczyk, Alasdair D.J. Freeman, Faith Leow, Timothy Wilson, David S. Rueda, David M.J. Lilley

## Abstract

Covalent linkages between chromosomes are naturally formed as a result of recombination or DNA replication. These can persist until late mitosis, resulting in interchromosomal ultrafine bridges, preventing cell division and causing genome instability. The endonuclease ANKLE1, localised at the cell midbody during cytokinesis and selective for DNA branchpoints, is ideally poised to act as an ‘enzyme of last resort’ to process interchromosomal bridges, enabling cell division to proceed. However, how ANKLE1 cleaves DNA under the tension existing in interchromosomal bridges during mitosis remains unexplored. Using optical tweezers, we show that ANKLE1 is a tension-stimulated endonuclease. High tension in DNA increases the ANKLE1 junction cleavage rate, with a twenty-fold increase at 60 pN. This indicates that ANKLE1 has evolved to respond to tension-induced DNA structural changes, thereby facilitating nucleolytic activity. This novel mechano-enzymological response of ANKLE1 reveals how it is well-suited to process ultrafine-bridges during late mitosis.

## Introduction

DNA junctions arise as a natural consequence of recombination and DNA replication, covalently linking chromosomes together. If incompletely processed, these junctions lead to the formation of ultrafine DNA bridges linking chromosomes during mitosis (1). Such DNA bridges can form in each cell cycle, their number increasing with DNA damage or inhibition of DNA replication (2). Interchromosomal bridges are thus a source of genome instability in cells—the persistence of a single linkage can prevent cell division or result in the mechanical rupture of DNA. Removing the DNA junctions that result in these bridges is so important that multiple pathways exist in eukaryotic cells for processing them. These include dissolution by BLM helicase and topoisomerase III or resolution by junction-selective nucleases, including the nuclear SLX4-SLX1-MUS81-EME1 complex and the cytoplasmic resolving enzyme GEN1 (reviewed in (3,4)). If junctions joining chromosomes have evaded processing by these pathways, one last enzyme can act late in the cell cycle to provide a final opportunity to process these remaining DNA branchpoints.

LEM-3 was first discovered as a GIY-YIG nuclease involved in DNA repair in *C. elegans* (5). Its mammalian (including human) ortholog, ANKLE1 (6,7), cleaves a variety of DNA junctions, including four-way Holliday junctions (4WJ), and three-way Y-junctions (i.e., a DNA duplex that separates into open single-stranded regions at one end) (8,9) (Figure 1A). The Y-junction was found to be the best substrate for human ANKLE1, cleaving three times faster than the four-way junction. Recent genetic and cytological evidence suggest that the biological role of LEM-3/ANKLE1 is processing ultrafine DNA bridges in anaphase. LEM-3/ANKLE1 has been shown to interact genetically with other junction-selective endonucleases, such as MUS81, GEN1, SLX1 and SLX4 (10). Furthermore, LEM-3/ANKLE1 accumulates at the cell mid-body (10,11), a structure where the two daughter cells abscise from each other. Genetic evidence indicates that such localisation is important for LEM-3/ANKLE1 activity *in vivo* (11). Finally, chromosomal bridges induced by treatment with DNA-damaging agents persist in *lem-3/ANKLE1* mutants (11). Thus, ANKLE1 is active at the right time, late mitosis, and present at the right location, the mid-body, to process remaining junctions. This suggests that ANKLE1 is the ‘enzyme of last resort’ in the processing of interchromosomal bridges to allow cell division to complete.

**Figure 1.**
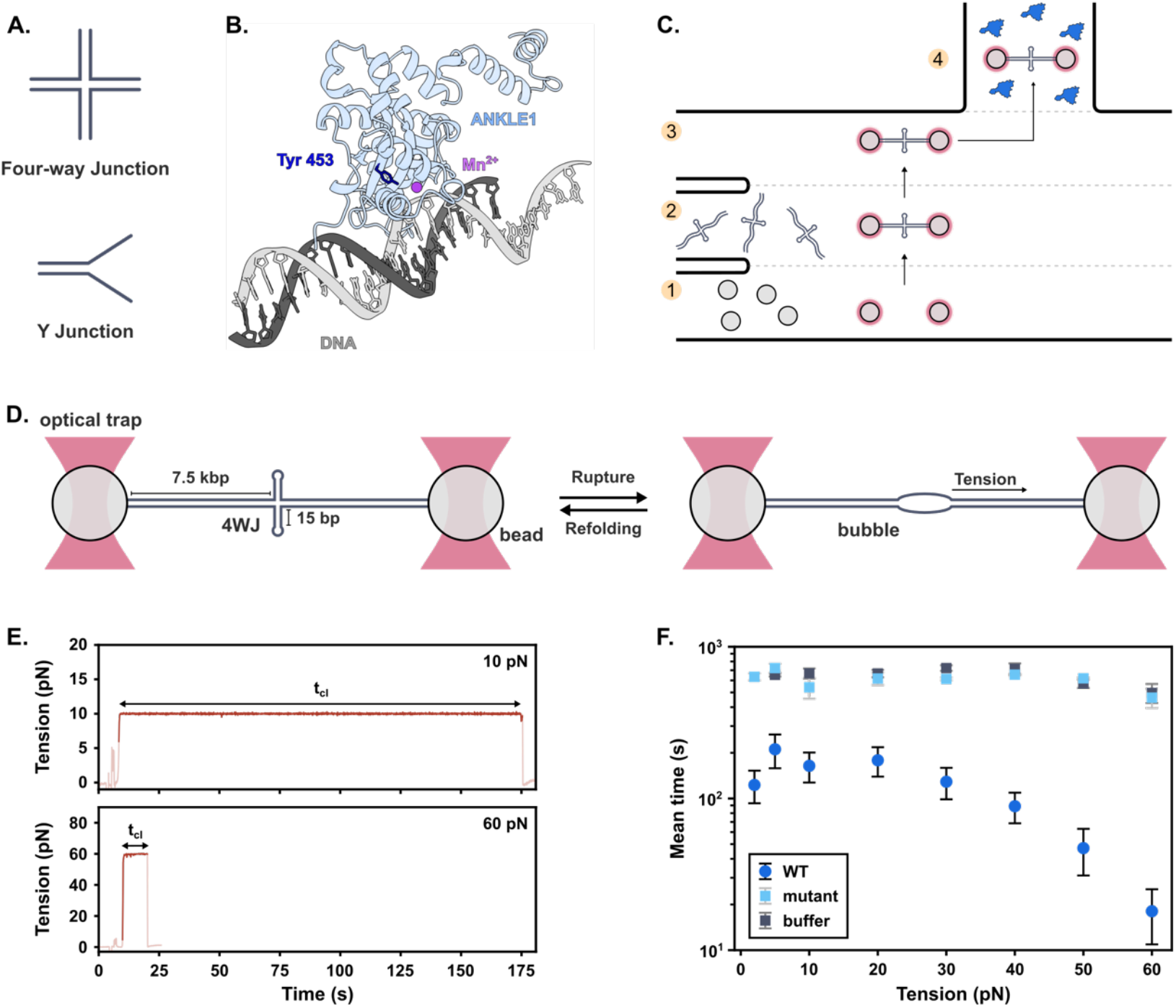
**A**) ANKLE1 substrates. **B)** Alphafold3-predicted structure of ANKLE1, highlighting Tyr453 (active-site), mutated in inactive ANKLE1 (9). **C)** Microfluidic chamber in optical tweezers with channels for (1) beads, (2) DNA, (3) buffer and (4) protein. **D)** 4WJ construct held between trapped beads showing 4WJ rupture. **E)** Representative tension-vs-time trajectories of 4WJ with ANKLE1 at low and high tension. Cleavage occurs where the tension falls to zero. **F)** Mean 4WJ cleavage time for WT ANKLE1 (N=78), inactive mutant (N=25) and buffer only (N=20). In the latter two, cleavage was sometimes not observed.

Segregation of chromosomes in anaphase requires pulling the unlinked chromosomes towards the spindle poles using pico-Newton (pN) mechanical forces generated by attached spindle fibres. The force generated to pull apart dividing chromosomes is estimated to be at least 50 pN (12,13). Any DNA bridging the separating chromosomes, thus must be under tension of a similar magnitude. To process these bridges, ANKLE1 must therefore act on DNA substrates under significant tension. Based on its suggested biological role, we hypothesised that the nuclease activity of ANKLE1 should respond to tension within its substrate. To test this, we used single-molecule optical tweezer experiments designed to study the ANKLE1-catalysed cleavage of DNA junctions (14) as a function of tension.

## Results and Discussion

We measured the cleavage rate of a 4WJ by ANKLE1 under tension using optical tweezers (14). We utilised the hANKLE_331–615 R492H_ nuclease domain of human ANKLE1 (referred to as ANKLE1) (Figure 1B) (9) and a 15 kbp central 4WJ construct with two 15 bp hairpins, as described (14), with individual DNAs tethered between two trapped beads. The 4WJ has been shown to rupture at ∼20-25pN into a bubble due to loss of base pairing of the helical arms (Figure 1C-D) (14). We observed a similar rupture under our conditions (Figure 2A), with a rupture force *F*_*rupture*_= 23 ± 2 pN (Figure 2B).

**Figure 2.**
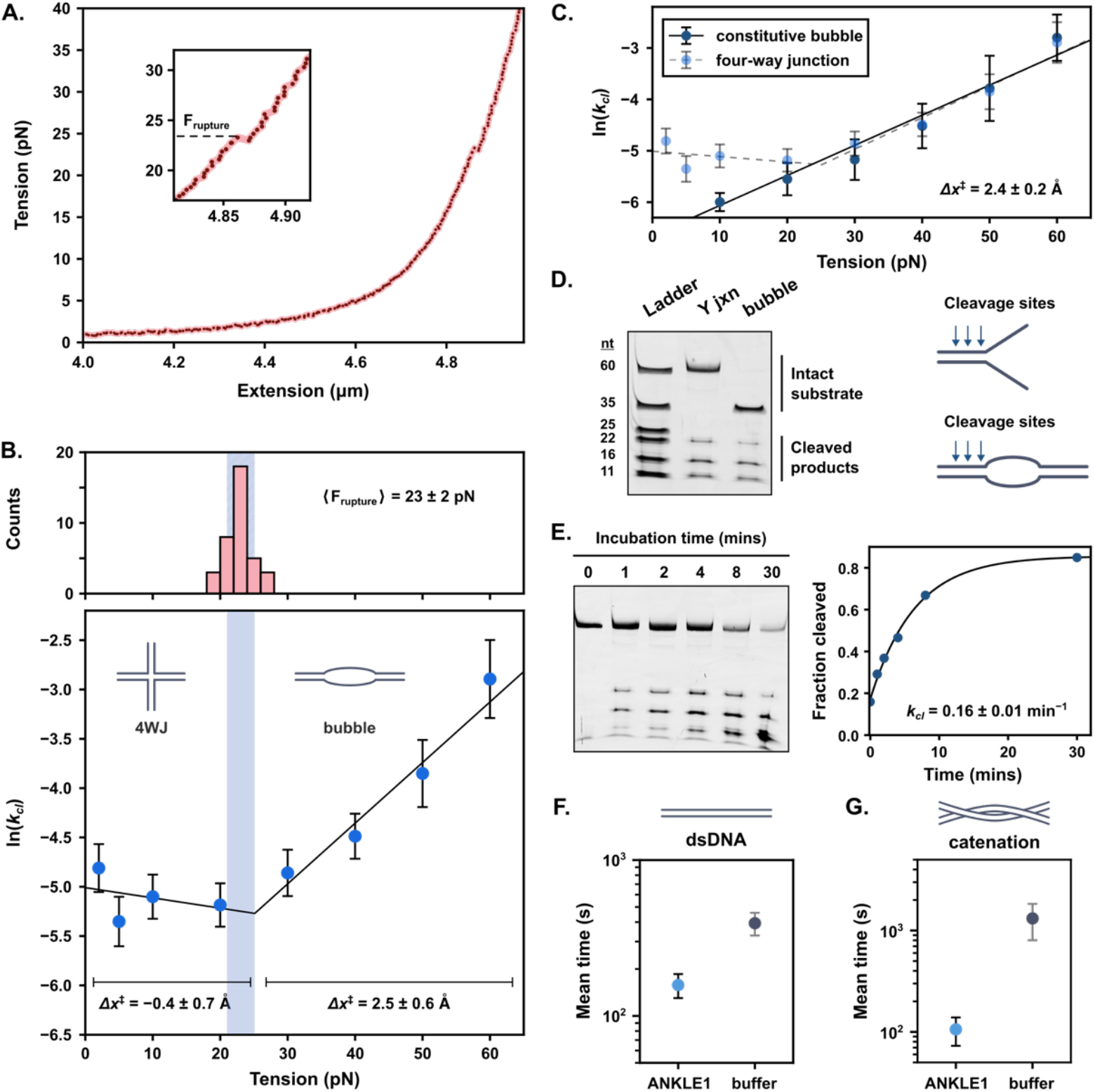
**A)** Representative force-extension plot of the 4WJ showing (inset) junction rupture at ∼23pN. **B)** Rupture-force histogram (top) and ln k_cl_ vs tension for the 4WJ (bottom) with Bell-Evans fits. Shaded regions correspond to 23 ± 2 pN. **C)** ln k_cl_ vs force for constitutive bubble (dark) and the 4WJ (light) with Bell-Evans fits. **D)** Denaturing polyacrylamide gel of substrates and products of nucleolytic cleavage for Y-junction (Y-jxn) and bubble substrates after 5min ANKLE1 incubation. Arrows (right) indicate ANKLE1 cleavage sites. **E)** Denaturing polyacrylamide gel of substrates and products of ANKLE1 cleavage as a function of time (left) and quantification (right) fitted to a single-exponential. **F)** Mean cleavage time of overstretched λ-DNA (60 pN) with ANKLE1 (N=25) or buffer (N=14). **G)** Mean cleavage time of catenated λ-DNA (40 pN) with ANKLE1 (N=5) or buffer (N=3).

Junction cleavage could be observed by holding the DNA at constant tension (force-clamp) and incubating with ANKLE1. The cleavage time (*t*_*cl*_) was measured from when tension was applied until it dropped to zero, indicating double-stranded DNA (dsDNA) cleavage (Figure 1E). We observed a marked dependence of *t*_*cl*_ on tension (Figure 1F), which was not observed in the absence of ANKLE1 or with a catalytically inactive mutant (ANKLE1_Y453F_) (9) (Figure 1F). Therefore, ANKLE1 is a tension-sensitive endonuclease.

According to the Bell-Evans model (15), the application of external force (F) modulates the free-energy barrier (Δ*G*^‡^), proportional to the logarithm of the cleavage rate (*k*_*cl*_= 1/⟨*t*_*cl*_⟩), by −*F*Δ*x*^‡^, where Δ*x*^‡^ represents the distance between the reactant state and the transition state along the reaction coordinate. We observed two distinct tension-dependent behaviours (Figure 2B). At low tension (<20-25 pN), In *k*_*cl*_ decreases slightly with tension, characterised by Δ*x*^‡^ = − 0.4 ± 0.7 Å, indicative of a weak force dependence. By contrast, at higher tension (>25 pN), we observed a larger positive force dependence, characterised by Δ*x*^‡^ = 2.5 ± 0.6 Å. At 60 pN, *k*_*cl*_ is ten-fold higher than at 20 pN. The inflection point between these behaviours occurs at 25 ± 5 pN, coinciding with the 4WJ rupture. This suggests that this inflection point arises from the 4WJ structure change, with the lower tension-dependence attributable to the intact 4WJ and the higher tension-dependence to the bubble (Figure 2B).

To test this, we utilised a construct containing two central 24nt DNA strands incapable of base-pairing with each other, forming a ‘constitutive’ bubble even at low forces. This construct exhibited a linear ln *k*_*cl*_ increase over the entire tension range, and a gradient corresponding to Δ*x*^‡^ = 2.4 ± 0.2 Å, nearly identical to the high-force regime for the four-way junction (Figure 2C). The rates of cleavage of the cruciform junction and bubble constructs are closely similar at tension above 25 pN (Figure 2C), indicating very similar activation parameters.

We expect the opening of the junction structure into an open bubble (Figure 1D) to create structures similar to Y-junctions (Figure 1A) at the bubble extremities. To test the equivalence of the Y-junction and bubble substrates for ANKLE1, we conducted cleavage experiments in bulk solution for both structures using short substrates with the same dsDNA sequences (see Materials and Methods). The bubble substrate is cleaved identically to the Y-junction (Figure 2D), with a *k*_*cl*_ = 0.16 min^−1^ (Figure 2E). Thus, our experiments indicate that ANKLE1 cleaves the Y-junction-like structures formed at the end of bubbles (i.e., the best substrate for this nuclease) upon rupture of junctions under tension. We expect these ruptured junctions to be present in ultrafine bridges during cytokinesis.

Previous studies have shown that bubbles containing stretches of single-stranded DNA can also be formed in overstretched dsDNA, in particular, around catenations (*i*.*e*., braids formed between two intertwined dsDNA molecules) (16). Such catenations form at the end of each round of DNA replication and can persist until anaphase, forming a separate, but equally prevalent class of ultrafine bridges (17). While not very active on dsDNA at zero force (8), ANKLE1 is capable of cleaving overstretched dsDNA held at forces above 60 pN (Figure 2F) and at DNA catenations held at high forces of around 40 pN (Figure 2G). This makes ANKLE1 an ideal endonuclease for the processing of all forms of anaphase bridges.

Finally, we see that the effect of tension on ANKLE1-mediated junction cleavage is not merely restricted to the formation of bubbles at elevated tension. If this were the case, we would expect the rate at which ANKLE1 cleaves junctions to plateau upon the formation of the bubble. Instead, *k*_*cl*_ increase with the applied tension well beyond the point we expect the junction to be disrupted, up to the tension believed to pull apart separating chromosomes. The sensitivity of ANKLE1 to tension is quite substantial. The rate of cleavage of the bubble increases 20-fold over the observed tension range.

We should emphasise that the observed change in rate with tension is that of a chemical reaction, the hydrolysis of phosphodiester bonds in the DNA. The origin of this rate acceleration is not known at present. The effect must result in an alteration in the reaction transition state that lowers the activation energy. Modulation of the active site structure likely results in ANKLE1 acting as a mechanosensitive endonuclease. It will be interesting to explore these changes in future studies.

## Conclusions

ANKLE1 has been proposed as the enzyme of last resort for the processing of any junction linking chromosomes. All its previously described characteristics point towards this role: it cleaves a variety of branchpoints in DNA, and it acts at the right time and location consistent with the cleavage of inter-chromosomal bridges. We further show that ANKLE1 preferentially cleaves bubble structures in DNA, likely generated by ruptured junctions or overstretched catenated DNA. Such structures are expected to be very common within ultrafine bridges. The activity of ANKLE1 is markedly stimulated by the level of tension expected in the bridges. Intriguingly, the topoisomerase Top2α, involved in decatenating DNA catenations under low tension, is inactivated at similar high tension (16). This is, therefore, further evidence for the proposed biological role of ANKLE1 in resolving ultrafine bridges and thus allowing cell division.

The mechano-enzymological response of ANKLE1 to tension within its DNA substrate is, to the best of our knowledge, the first example of a nuclease regulated by mechanical force on the DNA. But Nature tends to repeat the use of mechanisms, so this is now a property that might be sought in other enzymes operating on DNA.

## Materials and Methods

ANKLE1 was expressed in *E. coli* Arctic Express competent cells (Agilent) and purified using affinity and ion-exchange chromatography, as described (9). All optical tweezer experiments were performed as described (14).

Extended methods are provided in SI Appendix.

## Acknowledgments

We thank current and former members of the D.S.R. and D.M.J.L. labs for useful comments and discussions. D.S.R., K.K.R. and A.P.K. are supported by a core grant of the MRC-Laboratory of Medical Sciences (UKRI MC-A658-5TY10) and a Wellcome Trust Collaborative Grant (206292/Z/17/Z). D.M.J.L., A.D.J.F., F.L and T.W. were supported by a Cancer Research UK programme grant (A18604).

## Author Contributions

DSR and DMJL conceived initial studies. KKR, APK, ADJF, TW, DSR, and DMJL designed the experiments. KKR and APK performed optical tweezers experiments. KKR and DSR analysed the single-molecule data. ADJF and KKR purified the proteins. FL and TW prepared and purified the DNA. DMJL performed the bulk experiments. KKR, TW, DSR, and DMJL wrote the manuscript.

## Supporting Information for

### Extended Materials and Methods

#### Purification of ANKLE1

Wild-type ANKLE1 (hANKLE_331–615 R492H_) was expressed and purified as previously described (1). Briefly, a pGEX6P plasmid containing a GST-tagged ANKLE1 construct was transformed into *E. coli* Arctic Express competent cells (Agilent). The transformed cells were grown in LB containing 100 mg/l ampicillin and 20 mg/l gentamicin to an OD_600_ of 0.6– 0.8 at 37 ºC, then to an OD_600_ of 1.0–1.5 at 12 ºC. GST-ANKLE1 expression was subsequently induced with the addition of 200 µM IPTG at 12 ºC for 24 h. Cells were harvested by centrifugation at 7,500xg for 15 min at 4 ºC and resuspended in 40 ml of Lysis Buffer (25 mM HEPES pH = 7.5, 1 M NaCl, 5% glycerol, 0.5 mg/ml lysozyme, and 1 tablet of cOmplete EDTA-free protease inhibitor (Roche)) per litre of cell culture. The resuspended cells were incubated at room temperature for 15 min, supplemented with Triton X100 to a final concentration of 0.2% w/v, and lysed by sonication on ice for 1.5 min using 10 s pulses separated by 30 s. The lysate was clarified by centrifugation at 20,000xg for 30 min at 4 ºC and incubated with glutathione Sepharose 4B beads (Cytiva), which had been pre-equilibrated in GST Wash Buffer (25 mM HEPES pH = 7.5, 1 M NaCl, 5% glycerol). The beads were washed with GST Wash Buffer, and the GST tag was cleaved from the protein construct by incubation the beads with PreScission protease (GenScript) in GST Cleavage Buffer (25 mM HEPES pH = 7.5, 150 mM NaCl, 5% glycerol) overnight. The cleaved ANKLE1 was purified using two rounds of cation-exchange chromatography on a 5 ml HiTrap SP HP column (Cytiva) in 25 mM HEPES pH = 7.5, 10% glycerol with a linear gradient of 50 mM to 1 M NaCl and concentrated on a 1 ml HiTrap SP HP column (Cytiva) using a step-gradient up to 25 mM HEPES pH=7.5, 400 mM NaCl, 10% glycerol. Protein concentrations were measured using a Qubit 4 fluorometer and single-use aliquots were flash frozen in liquid nitrogen and stored at −80 ºC.

#### Preparation of 4WJ construct for optical tweezer experiments

The custom 15 kb dsDNA construct containing a centrally located four-way junction was prepared as previously described (2). Briefly, a 7.5 kb handle was prepared using PCR with Pfu Ultra II Fusion HS polymerase (Agilent) from a *λ* -phage genomic DNA template using two modified primers, Biot-F (which contained four biotins in the 5’ end) and AS-R2 (which contained a single abasic site). The sequences of these primers are given below. The central four-way junction was formed by annealing two oligonucleotides, HX-J3a and HX-J3b (sequences given below) by slow-cooling an equimolar mixture of the two from 80 ºC to room temperature in Hybridisation Buffer (10 mM Tris-HCl pH_RT_ = 7.5, 50 mM NaCl). The annealed junction ligated to the handles on both ends at 16 ºC for 8 h using T4 DNA ligase (New England Biolabs) and purified by electrophoresis in a 0.6% agarose, 1x TAE gel and electroelution of the product band.

The oligonucleotide sequences are:

**Biot-F**: (5’-Biot)-(Biot-dT)CA(Biot-dT)C(Biot-dT)GAAA CAGCAGCGGA,

**AS-R2**: **p**CTTACAGATG (ab)CGCACGAAA AGCATCAGGT C,

**HX-J3a**: **p**ACATCTGTAA GAGTCTGCAG TTGAGTCCTT GCTAGGACGG ACGAAGTC-CG TCCTAGCAAG GGGCTGCTAC CGGAAG,

**HX-J3b**: **p**ACATCTGTAA GCTTCCGGTA GCAGCCTGA GCGGTG GATG AACGAAGT-TC ATCCACCGCT CAACTCAACT GCAGACT,

where (5’-Biot) denotes 5’-biotin, (Biot-dT) denotes biotin-dT, (ab) denotes the abasic dSpacer CE, and **p** denotes 5’ phosphorylation.

#### Preparation of bubble construct for optical tweezer experiments

The custom 15 kb bubble construct consisted of two 7.5 kb dsDNA handles and a centrally located constitutive ‘bubble’, *i*.*e*., a permanent, unpaired region of 24 nt. The 7.5 kb handles were prepared in a manner analogous to the ones for the 4WJ construct by PCR amplification from a λ -phage genomic DNA template using the Biot-F primer (see above) and a second modified primer, Fluor-R. Fluor-R contained a *BsaI* recognition site which could be used to generate an asymmetric 4 nt 5’ overhang in the handle for ligation to the central bubble and 5’-fluorescein label that can be used to follow the generation of this overhang (sequence given below). The primers were synthesised on an Applied Biosystems 394 synthesiser using β -cyanoethyl phosphoramidites (Glen Research) (3, 4), deprotected by incubation with ammonium hydroxide at 55 ºC for 2 h, and purified by electrophoresis in polyacrylamide gels under denaturing conditions in 7 M urea. The primers were recovered from DNA-containing gel fragments using the crush and soak method followed by ethanol precipitation. PCR amplification of the 7.5 kb handles from a λ -phage genomic DNA template was performed using the above primers and Q5 High-Fidelity Master Mix (New England Biolabs) using the manufacturer’s recommended cycling protocol, and purified using the QIAquick PCR purification kit (Qiagen).

The central bubble was generated from two chemically synthesised phosphorylated oligonucleotides (Thermo Scientific), denoted bub-a and bub-b (sequences given below). The oligonucleotides were purified by electrophoresis in polyacrylamide gels under denaturing conditions in 7 M urea and recovered from DNA-containing gel fragments using the crush and soak method followed by ethanol precipitation. The bubble was formed by mixing equimolar amounts of both oligonucleotides in Hybridisation Buffer supplemented with 50 mM NaOAc, heating to 80 ºC, and slowly cooling to room temperature. This generated an insert with 4 nt 5’-GGTT overhangs at each end that were complementary to the 5’-AACC overhangs generated by *Bsa*I digestion of the DNA handles, but not to themselves, thereby preventing self-ligation. The reaction was cycled in a thermocycler (Techne TC-3000G) for 30 cycles of 5 min at 37 °C (for *Bsa*I digestion) followed by 5 min at 16 °C (for ligation) and terminated by purifying the product using the QIAquick PCR purification kit (Qiagen). The ligated 15 kb construct was further purified by electrophoresis in a 0.8% agarose gel prepared in TAE buffer and stained with 0.5x SYBR Safe DNA Gel Stain (Invitrogen). The product band was excised, and DNA was recovered using the QIAEX II Gel Extraction Kit (Qiagen).

The oligonucleotide sequences are:

**Fluor-R:** (5′-Fluor)-CCGTTGC**GGT CTC**AAACCGC ACGAAAAGCA TCAGGTC,

**bub-a: p**GGTTGCAGTT GAGTCCTTGC TAGGACGGAT CCCTCGAGGC TAGGTACCT,

**bub-b: p**GGTTAGGTAC CTAGCATCTG TGAATTCAAC CACCGCTCAA CTCAACTGC,

where (5′-Fluor) denotes 5’-fluorescein, the nucleotides in bold denote the *BsaI* recognition site, and **p** denotes 5’ phosphorylation.

#### Preparation of λDNA construct for optical tweezers experiments

The 48.5 kb λ-DNA was prepared as previously described (5). Briefly, 4 nM λ-phage genomic DNA (Thermo Scientific) was incubated with 100 µM dGTP, 100 µM dTTP, 80 µM biotin-14-dATP, 80 µM biotin-14-dCTP, and 0.5 U Klenow polymerase exo^−^ (New England Biolabs) in NEB2 Buffer (New England Biolabs) at 37 ºC for 30 min, then at 70 ºC for 15 min, before being purified using the QIAquick PCR purification kit (Qiagen).

#### Optical tweezer experiments

Single-molecule ANKLE1 cleavage experiments were performed on a C-trap (Lumicks), integrating optical tweezers, microfluidics, and confocal fluorescence microscopy. Prior to the experiments, all solutions were filtered using 0.2 µm syringe filters. The microfluidic flowcell and channels were passivated with 0.5% w/v Pluronic F-127 in TN50 Buffer (50 mM Tris-HCl pH_RT_ = 7.5, 50 mM NaCl) and subsequently, for channels 3 and 4 (Figure 1), with ANKLE1 Experiment Buffer (50 mM Tris-HCl pH_RT_ = 7.5, 40 mM NaCl, 10 mM MnCl_2_, 1 mg/ml BSA). At the start of the experiment, the following substrates were introduced into the corresponding channels by flow (Figure 1): (i) 0.005% w/v streptavidin-coated polystyrene beads (∼4.4 µm, Spherotech) in TN50 Buffer in channel 1; (ii) ∼2 pM of biotinylated DNA substrate (4WJ or bubble or λ DNA construct) in TN50 Buffer in channel 2; (iii) ANKLE1 Experiment Buffer in channel 3; (iv) 500nM ANKLE1 (wild-type or mutant) in ANKLE1 Experiment Buffer in channel 4. The optical trap stiffness used in these experiments was 0.2–0.3 pN/nm. For the dual-trap experiments, two beads were optically trapped in channel 1 and then moved to channel 2, where a single biotinylated DNA molecule was tethered between them. The presence of the correct DNA tether was verified by moving the DNA to channel 3 and generating a force-extension curve. For the 4WJ construct, only DNA molecules, which showed a rupture of the junction, were kept for further experiments. The DNA was subsequently moved to the protein channel in a relaxed state (at an extension of ∼2.5 µm for the 15 kb constructs, or of ∼8 µm for the 48.5 kb λ DNA construct) and a force clamp was applied to stretch the DNA to the desired tension. The tension on the DNA and the corresponding distance were recorded from before the start of the application of the force clamp till cleavage of the DNA was observed (or in case of incubations with mutant ANKLE1 or just ANKLE1 Experiment Buffer, for ∼10 min if no cleavage was observed till then).

For quadruple trap experiments, a single λ DNA molecule was tethered between each of two pairs of optically trapped beads and verified using force-extension curves. The two DNA molecules were subsequently braided once to form a single right-handed catenation, in a manner similar to (6), by moving one pair of beads above, around, and then below the other pair (Supplementary Movie S1). Establishment of the catenation was verified by measuring simultaneous increase in the force measured on both pair of beads when only one pair was moved—which would only be possible if there was a braid formed between the DNA molecules tethered to both pairs of beads. The beads were then moved into the protein channel with the DNA molecules in a relaxed position and one of the beads was moved till a high force (∼40 pN) was measured for that bead pair. The forces on both pairs of beads and the corresponding distances were recorded till a simultaneous reduction in the forces (to zero for one bead pair and a low, non-zero value for the other) was observed, signifying cleavage on one of the two intertwined DNA molecules.

#### Single-molecule data analysis

All analyses of the single-molecule data generated above were performed using Python code on Jupyter notebooks. The software package pylake (Lumicks) was used for handling the raw data generated from the C-trap experiments. For the DNA cleavage assays, the application of the force clamp and the cleavage of DNA were detected using a threshold force lower than the force applied in the clamp. The activation of the clamp was detected when the measured force crossed the threshold from below and DNA cleavage was detected when the force crossed the threshold from above. The time between these two events was denoted as the time to cleavage (*t*_*cl*_). For instances (in the case of incubation with mutant ANKLE1 or just ANKLE1 Experiment Buffer) where no cleavage was observed, the end of the force recording was chosen instead. Since the cleavage of DNA occurred within a single time-point and the force-clamp achieved the desired force rapidly, the specific choice of the threshold force did not affect the calculated *t*_*cl*_ by more than a few milliseconds.

For the 4WJ and bubble constructs, the average time to cleavage (⟨*t*_*cl*_ ⟩) was converted to the rate of cleavage (*k*_*cl*_) using the relation

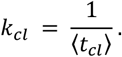

For the 4WJ construct, *l*n *k*_*cl*_ was fit to a piecewise linear function given by

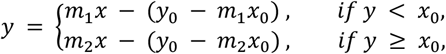

where *x*_0_is the inflection point, *y*_0_ is the value of the function at *x*_0_, and *m*_1_and *m*_*2*_are the slopes before and after *x*_0_ respectively. The inflection point *x*_0_ was a free parameter and determined from the fit.

For the bubble construct, In *k*_*cl*_ was fit to a linear function given by

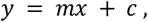

where *m* is the slope of the line and *c* is the intercept.

According to the Bell-Evans model (7), at any given force *F*,

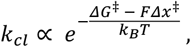

where *ΔG*^‡^ is the activation free energy at zero force, *Δx*^‡^ is the distance along the relevant reaction coordinate between the reactant ground state and the transition state, *k*_*B*_ is the Boltzmann constant, and *T* is the absolute temperature (taken to be 298 K for these experiments). Using the above relation, the slopes of the fits for *l*n *k*_*cl*_ *versus F* were converted to the corresponding *Δx*^‡^ using

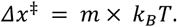

#### Bulk ANKLE1 cleavage experiments

The 5’-fluorescein-labelled short Y junction construct (sequence given below) used for bulk cleavage was prepared as previously described (1). The 5’-fluorescein-labelled short bubble construct (sequence given below) was prepared in a manner analogous to the central bubble insert described above. As previously described (1), 50 nM fluorescein-labelled DNA was incubated with 2 µM ANKLE1 for 3 min at 37°C in 20 mM cacodylate (pH 6.5), 50 mM KCl. The DNA cleavage reaction was then initiated by addition of 10 mM MnCl_2_. Aliquots were removed at chosen times, and the reaction was terminated by addition of two volumes of formamide. Substrate and cleavage products were separated by gel electrophoresis in 17% polyacrylamide gels under denaturing conditions and imaged using a Typhoon FLA 9500 fluorimager (Fuji). The data were quantified in Image Guage 4.21 (Fuji) and fit to a single-exponential function of the form

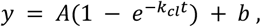

where *k*_*cl*_ is the bulk rate of cleavage, *a* is the amplitude of the reaction, and *b* is the background.

The oligonucleotide sequences are:

**YJXa:** (5’ Fluor)-CATGTGTTCT AGACTGCAGT TGAGTTCTAG CTTCT

**YJXb:** TCGTCTGACT ACTCAACTGC AGTCTAGAAC ACATG

**bub2-a:** (5’-Fluor)-CATGTGTTCT AGACTGCAGT TGAGTTCTAG CTTCTCCTTG CTAG-GACGGA TCCCTCGAGG

**bub2-b:** CATGTGTTCT AGACTGCAGT TGAGTTCTAG CTTCTCCTTG CTAGGACGGA TCCCTCGAGG

where (5′-Fluor) denotes 5’-fluorescein.

### Supplementary movie

#### Movie S1 (separate file)

Video showing two molecules of λ DNA tethered between four optically trapped beads in the quadruple-trap setup. Beads 1 and 2 are moved over, around, and then under beads 3 and 4, to form a right-handed catenation between the two DNA molecules.

## References

1. K. L. Chan, I. D. Hickson, New insights into the formation and resolution of ultra-fine anaphase bridges. Semin. Cell Dev. Biol. 22, 906–912 (2011).

2. Y. W. Chan, K. Fugger, S. C. West, Unresolved recombination intermediates lead to ultra-fine anaphase bridges, chromosome breaks and aberrations. Nat. Cell. Biol. 20, 92–103 (2018).

3. Y. Liu, A. Freeman, A.-C. Déclais, A. Gartner, D. M. J. Lilley, Biochemical and Structural Properties of Fungal Holliday Junction-Resolving Enzymes. Methods Enzymol. 543–568 (2018).

4. S. Sarbajna, S. C. West, Holliday junction processing enzymes as guardians of genome stability. Trends Biochem. Sci. 39, 409–419 (2014).

5. C. M. Dittrich, et al., LEM-3 – A LEM Domain Containing Nuclease Involved in the DNA Damage Response in C. elegans. PLoS One 7, e24555 (2012).

6. A. Brachner, et al., The endonuclease Ankle1 requires its LEM and GIY-YIG motifs for DNA cleavage in vivo. J. Cell Sci. 125, 1048–1057 (2012).

7. J. Braun, A. Meixner, A. Brachner, R. Foisner, The GIY-YIG Type Endonuclease Ankyrin Repeat and LEM Domain-Containing Protein 1 (ANKLE1) Is Dispensable for Mouse Hematopoiesis. PLoS One 11, e0152278 (2016).

8. J. Song, A. D. J. Freeman, A. Knebel, A. Gartner, D. M. J. Lilley, Human ANKLE1 Is a Nuclease Specific for Branched DNA. J. Mol. Biol. 432, 5825–5834 (2020).

9. A. D. J. Freeman, A.-C. Déclais, T. J. Wilson, D. M. J. Lilley, Biochemical and mechanistic analysis of the cleavage of branched DNA by human ANKLE1. Nucleic Acids Res. 51, 5743–5754 (2023).

10. H. Jiang, N. Kong, Z. Liu, S. C. West, Y. W. Chan, Human Endonuclease ANKLE1 Localizes at the Midbody and Processes Chromatin Bridges to Prevent DNA Damage and cGAS-STING Activation. Adv. Sci. 10, 2204388 (2023).

11. Y. Hong, et al., LEM-3 is a midbody-tethered DNA nuclease that resolves chromatin bridges during late mitosis. Nat. Commun. 9, 728 (2018).

12. E. L. Grishchuk, M. I. Molodtsov, F. I. Ataullakhanov, J. R. McIntosh, Force production by disassembling microtubules. Nature 438, 384–388 (2005).

13. F. Rago, I. M. Cheeseman, The functions and consequences of force at kinetochores. J. Cell Biol. 200, 557–565 (2013).

14. A. P. Kaczmarczyk, et al., Search and processing of Holliday junctions within long DNA by junction-resolving enzymes. Nat. Commun. 13, 5921 (2022).

15. E. A. Evans, D. A. Calderwood, Forces and Bond Dynamics in Cell Adhesion. Science 316, 1148–1153 (2007).

16. E. E. Cutts, S. Saravanan, G. L. M. Fisher, D. S. Rueda, L. Aragon, Substrate accessibility regulation of human TopIIa decatenation by cohesin. [Preprint] (2023). Available at: https://www.biorxiv.org/content/10.1101/2023.11.20.567865v1.

17. A. Finardi, L. F. Massari, R. Visintin, Anaphase Bridges: Not All Natural Fibers Are Healthy. Genes 11, 902 (2020).

## SI References

1. A. D. J. Freeman, A.-C. Déclais, T. J. Wilson, D. M. J. Lilley, Biochemical and mechanistic analysis of the cleavage of branched DNA by human ANKLE1. Nucleic Acids Res. 51, 5743–5754 (2023).

2. A. P. Kaczmarczyk, et al., Search and processing of Holliday junctions within long DNA by junction-resolving enzymes. Nat. Commun. 13, 5921 (2022).

3. S. L. Beaucage, M. H. Caruthers, Deoxynucleoside phosphoramidites—A new class of key intermediates for deoxypolynucleotide synthesis. Tetrahedron Lett. 22, 1859–1862 (1981).

4. N. D. Sinha, J. Biernat, J. McManus, H. Köster, Polymer support oligonucleotide synthesis XVIII: use of β-cyanoethyi-N,N-dialkylamino-/N-morpholino phosphoramidite of deoxynucleosides for the synthesis of DNA fragments simplifying deprotection and isolation of the final product. Nucleic Acids Res. 12, 4539–4557 (1984).

5. P. Gross, G. Farge, E. J. G. Peterman, G. J. L. Wuite, Combining Optical Tweezers, Single-Molecule Fluorescence Microscopy, and Microfluidics for Studies of DNA–Protein Interactions. Methods Enzymol. 475, 427–453 (2010).

6. E. E. Cutts, S. Saravanan, G. L. M. Fisher, D. S. Rueda, L. Aragon, Substrate accessibility regulation of human TopIIa decatenation by cohesin. [Preprint] (2023). Available at: https://www.biorxiv.org/content/10.1101/2023.11.20.567865v1.

7. E. A. Evans, D. A. Calderwood, Forces and Bond Dynamics in Cell Adhesion. Science 316, 1148–1153 (2007).

